# Global Pannexin 1-deletion increases tumor-infiltrating lymphocytes in the BRAF/Pten mouse melanoma model

**DOI:** 10.1101/2023.07.28.551021

**Authors:** Rafael E. Sanchez-Pupo, Garth A. Finch, Danielle E. Johnston, Heather Craig, Rober Abdo, Kevin Barr, Steven Kerfoot, Lina Dagnino, Silvia Penuela

## Abstract

Immunotherapies for malignant melanoma seek to boost the anti-tumoral response of CD8^+^ T cells but have a limited patient response rate, in part due to limited tumoral immune cell infiltration. Genetic or pharmacological inhibition of Pannexin 1 (PANX1) channel-forming protein is known to decrease melanoma cell tumorigenic properties *in vitro* and *ex vivo*. Here, we crossed *Panx1* knockout (*Panx1*^−/−^) mice with the inducible melanoma model: *Braf*^CA^, *Pten*^loxP^, *Tyr::CreER^T2^* (BPC). We found that deleting the *Panx1* gene in mice does not reduce BRAF(V600E)/Pten-driven primary tumor formation or improve survival. However, BPC-*Panx1*^−/-^ mice tumors exhibited a significant increase in infiltration of CD8^+^ T lymphocytes with no changes in the expression of early T cell activation marker CD69, LAG-3 checkpoint receptor or PD-L1 in tumors when compared to BPC-*Panx1*^+/+^ genotype. Our results suggest that although a *Panx1* deletion does not overturn the aggressive *BRAF*/*Pten*-driven melanoma progression *in vivo*, it does increase the infiltration of effector immune T cell populations in the tumor microenvironment. We propose that PANX1-targeted therapy could be explored as a strategy to increase tumor-infiltrating lymphocytes to boost anti-tumor immunity.

## INTRODUCTION

Malignant melanoma is a type of skin cancer that develops from melanocytes and becomes a life-threatening disease when it spreads in the body [1]. Melanomas can originate from diverse primary sites (e.g., cutaneous, ocular, or mucosal), however, approximately 3% of the cases exhibit a lack of an identifiable primary site (known as: melanoma of unknown primary - MUP), which have been less understood in terms of their biological characteristics when compared to the more conventional melanomas of known primary (also known as MKP) [2]. While the prognostic factors for MUP seems to be similar to MKP at the same stage, patients with MUP tends to experience better outcomes in comparison to the latter group. This improvement in prognosis has been attributed to higher immunogenicity, which has been evident in the immune-mediated regression of the primary tumor site [2].

Depending on the diagnosis and the mutational landscape of the tumor biopsies (e.g., *BRAF* mutation status), targeted therapy with mutant kinase inhibitors or immunotherapies with monoclonal antibodies against immune checkpoint inhibitors (e.g., anti-CTLA4, anti-PD-1 antibodies) are currently being used for melanoma treatment [3]. As research into the PI3K (Phosphoinositide 3-kinase)/AKT (Protein kinase B) /NF-κB pathway deepens, its role in melanoma has also attracted significant attention. Dysregulating mutations of AKT family members is a common occurrence in melanoma, being identified in as much as 43-60% of melanoma cases [4]. PTEN, a protein phosphatase that typically restrains the cell-growth signaling cascade of PI3K/AKT/mTOR, has been found to be altered in 38% of primary melanoma patients and 58% of those with metastatic disease [5]. Notably, alterations in both PTEN and the BRAF genes also coexist, potentially leading to the simultaneous dysregulation of both the Mitogen-activated protein kinases (MAPK) and PI3K pathways. Consequently, there is a possibility that PI3K inhibitors may offer some therapeutic benefit to patients with PTEN and/or AKT-mutant melanomas [6]. On the other hand, the use of immune checkpoint inhibitors enhances anti-tumoral T cell-mediated immune responses, but unfortunately, about ∼60% of melanoma patients do not respond to this type of therapy [7–9]. Regardless of the treatment options, their effectiveness is usually dampened due to the insufficient effector activity of tumor-infiltrating lymphocytes (TILs) [10], recurrence, and development of multiple resistance mechanisms [11, 12]; therefore, innovative therapy concepts are still warranted.

Pannexin1 (PANX1) is a channel-forming membrane glycoprotein that mediates the passage of nucleotides, ions, and other metabolites involved in intercellular communication [13]. We have previously demonstrated that PANX1 expression correlates with tumor cell aggressiveness in isogenic mouse melanoma cell lines, and its knockdown reduces melanoma tumor size and metastasis to the liver in chick chorioallantoic membrane (CAM) xenografts [14]. More recently, we showed that PANX1 is highly expressed in human melanomas, and genetic or pharmacological targeting of PANX1 decreases the tumorigenic properties of melanoma cells [15]. Mechanistically, PANX1 regulates melanoma cell growth and metabolism through direct interaction with β-catenin and modulation of the Wnt signaling pathway [16]. All these findings place PANX1 as an attractive target for preclinical melanoma research, but it is still unclear whether the silencing of this gene would impact melanoma development *in vivo*.

In addition, PANX1 influences the inflammatory response [17–22] and is expressed in different subtypes of immune cells like macrophages and lymphocytes [17, 23, 24]. Previous evidence shows that PANX1-mediated adenosine-5′-triphosphate (ATP) permeability enables activation and functional enhancement of T cells [21, 22]. On the other hand, the release of ATP through caspase-cleaved PANX1 channels in apoptotic leukemic lymphocytes has been shown to induce the recruitment of inflammatory cells (e.g., monocytes)[17]. Paradoxically, the caspase-dependent PANX1 channel opening in dying macrophages and lymphocytes releases metabolites that suppress, *in vivo,* proinflammatory signals of myeloid cells in the surrounding tissue [13]. Considering the broad PANX1 expression and different cell types present in the tumor microenvironment (TME), no studies have specifically addressed whether PANX1 modulates the recruitment of immune cells in the melanoma TME [18].

Previous histological and pathological examinations of a constitutive global *Panx1* knockout mouse (*Panx1*^−/-^, *Panx1*KO)(developed by Genentech) [20] showed no overt phenotypes. However, using the same *Panx1*^−/-^ mouse strain, we have found differences in dorsal skin thickness and delayed wound healing [14, 25]. Moreover, our studies show that in *Panx1* wildtype mice, normal melanocytes express low levels of PANX1 which are upregulated in melanoma cells [14]. To gain new insights into the roles of PANX1 in the context of skin cancer we set out to test the combination of this *Panx1*^−/-^ mouse line with a cutaneous melanoma mouse model.

In this work, we crossed the global *Panx1*^−/-^ mouse strain with the inducible melanoma model *BRaf^CA/+^, Pten^LoxP/Loxp^, Tyr::CreER^T2^* (abbreviated here as BPC) that harbors two known oncogenic driver mutations [26–28]. Due to the role of PANX1 in regulating tumor growth and the immune response [17–22], we sought to evaluate the effects of the *Panx1* germline deletion on melanoma progression and the tumor immune infiltration of this new hybrid mouse model. Our results showed that *Panx1* global deletion did not reduce the strong BRAF/Pten-driven melanoma progression but increased the tumor infiltration of effector immune T cell populations. Interestingly, PANX1-deficient BPC mice exhibited sex-driven morphological differences in spleen size without an apparent influence on the tumor burden. We anticipate that targeting PANX1 in melanoma may increase the homing of effector T-lymphocytes to melanoma tumors which in the future could be harnessed to improve the effect of immunotherapies.

## Materials and Methods

### Mouse breeding and genotyping

C57BL/6, Tyr::CreER^T2^; Braf^CA/+^; Pten^LoxP/LoxP^ (BPC) mice were obtained from Jackson Laboratory, (Stock # 013590). Mice were kept on a 12:12 h light:dark cycle with normal chow and water available *ab libitum*. For deletion of *Panx1*, BPC mice were crossed with the global *Panx1*^−/−^ mice in a C57BL/6N background strain [25, 20] (Genentech Inc., San Francisco, CA). Genotyping was performed on genomic DNA derived from the toes using PCR as described in [27].

### Melanoma tumor induction, monitoring and sample collection

All experiments performed on the mice were approved by the Animal Care Committee of the University Council on Animal Care at the University of Western Ontario, London ON, Canada (UWO AUP# 2019-070). 3–4-weeks-old BPC mice were used for tumor induction experiments: *Panx1*^+/+^ (7 females and 8 males) and *Panx1*^−/−^ (9 females and 4 males). Cutaneous melanomas were induced with topical application of 4-hydroxytamoxifen (4-HT) (Sigma) on the skin of the lower back as per protocol in [27]. Mouse body weight and tumor dimensions were monitored every 3–4 days as soon as the tumors were visible. Measurements were done with a digital caliper, and tumor volume was calculated using the modified ellipsoid formula V = (W(×2) × L)/2, where V is tumor volume, L and W are the longest (length) and shortest (width) tumor diameters, respectively. The endpoint was considered when the tumor volume (or total combined tumor volume, in case of multiple tumors per animal) reached 2.0 cm^3^, tumors showed signs of heavy ulceration, or the animals had poor body condition. Euthanasia was carried out using carbon dioxide asphyxiation followed by cervical dislocation. The spleens were weighed and measured using a ruler. Draining inguinal lymph nodes compromised with pigmented lesions were classified as metastatic sites and counted per mouse. Tissue samples were fixed overnight with Periodate-Lysine-Paraformaldehyde (PLP) fixative or flash-frozen in liquid nitrogen and stored at –80 °C until processing for RNA extraction.

### Tissue Processing, RNA Extraction, real time-qPCR

Frozen tissues/tumor samples were ground in liquid nitrogen and total RNA was extracted using TRIzol™ Reagent (Invitrogen) according to the manufacturer’s protocol and purified using the RNAeasy® Plus Mini kit (Qiagen). RNA concentration was determined using an Epoch Microplate Spectrophotometer (BioTek®). cDNA was generated using the High-Capacity cDNA Synthesis Kit (Thermo Fisher Scientific; REF 4368814) in a T100^TM^ Thermal Cycler (Bio-Rad). SYBR®Green Real-Time PCR Master Mix (Bio-Rad) was used along with the primers listed in **Table 1**, where mouse *gapdh* and *18S* were the housekeeping gene controls. Real-time qPCR was performed in a CFX96 Touch TM Real-Time PCR Detection System (Bio-Rad). All the assays involved at least three technical replicates (n=3), and the relative mRNA expression analysis was determined using the ΔΔCt method calculated in the Bio-rad CFX Maestro Software, ver 1.1 (Bio-rad). Between assays, a non-tamoxifen-treated skin BPC-*Panx1*^+/+^ sample was used as the control for inter-plate variation.

**Table 1.**
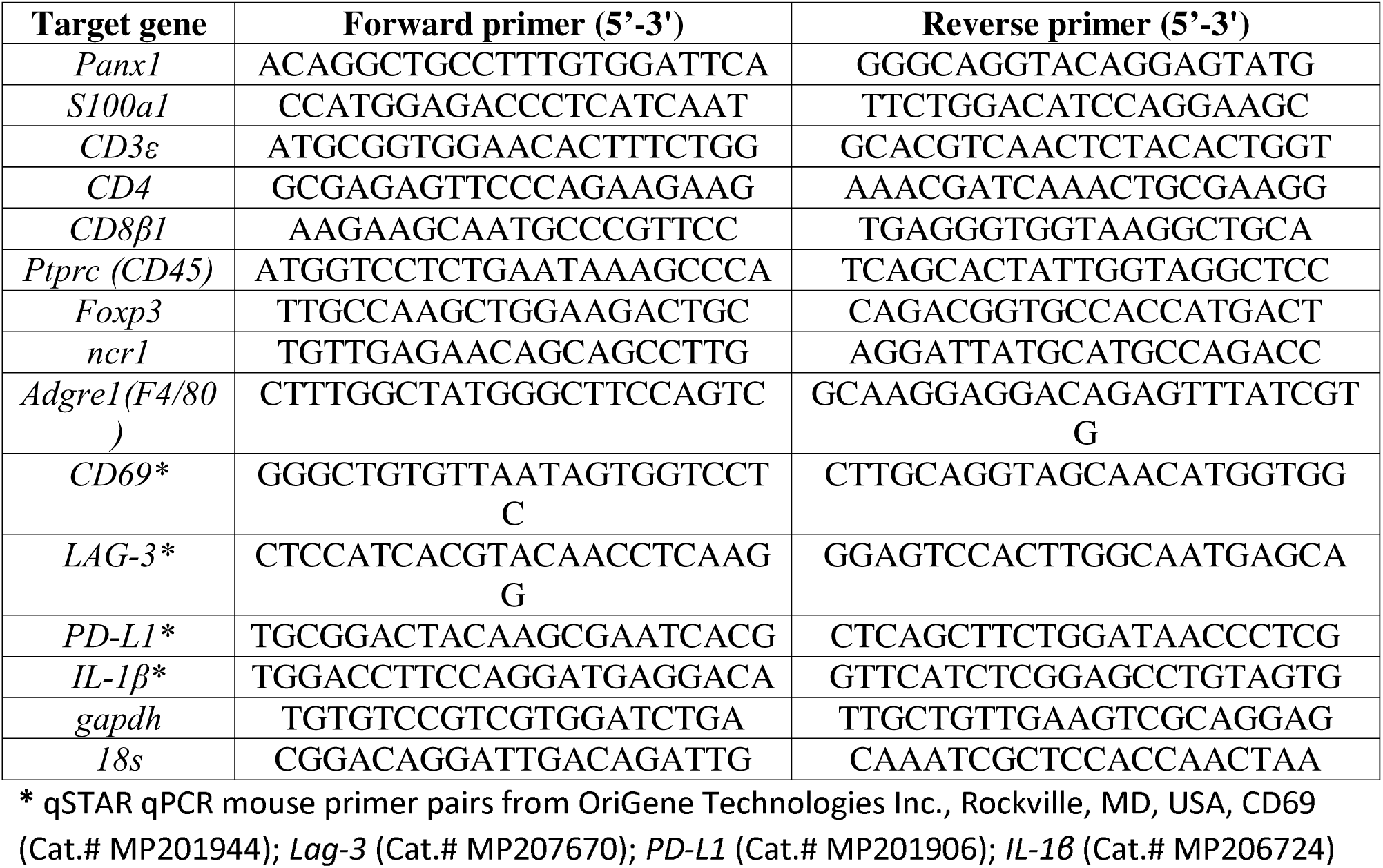
List of primers used for real-time qPCR analysis.

### Tissue preparation, cryo-sectioning, and immunofluorescence staining

Tumor tissues samples that were fixed in PLP (1% PFA, 0.1 M L-Lysine, 0.01 M sodium periodate) overnight at 4°C were washed three times for 5 mins in phosphate buffer (0.075 M sodium phosphate dibasic and 0.025 M sodium phosphate monobasic) and incubated sequentially in 10%, 20%, and 30% sucrose phosphate solutions, each overnight at 4°C. Tumors were then embedded and frozen in Fisher Healthcare™ Tissue-Plus™ O.C.T. Compound (Fisher Scientific Cat.# 23-730-571) and stored at -80 °C until sectioning. Eight-micron-thick serial cryostat sections were immuno-stained with the fluorescent-conjugated antibodies diluted in 1% BSA, 0.1% Tween-20, and 5% rat-serum blocking buffer. Counterstaining was done with 10 µg/mL of Hoechst 33342 (Life Technologies Ref.# H3570) for 7 min, at room temperature. Sections were mounted with ProLong™ Gold Antifade Mountant (ThermoFisher Cat. # P10144).

### Antibodies used for immunofluorescence

Alexa Fluor® 647-conjugated anti-mouse CD4 (clone RM4-5) and anti-mouse CD8a (clone 53-6.7), (BioLegend, USA, Cat.#: 100530 and 100727, respectively) were used for immunofluorescence at a final concentration of 2.5 µg/mL and eFluor 660 anti-mouse FOXP3 (clone FJK-16s) eBioscience™ (ThermoFisher Scientific, Cat.# 50-5773-80) was used at 1 µg/mL. eFluor 660 rat IgG2a kappa Isotype Control (eBR2a) (eBioscience™, ThermoFisher Sci., Cat. # 50-4321-80) and Alexa Fluor® 647 Rat IgG2a, κ Isotype Ctrl (BioLegend, USA, Cat. #: 400526) were used as controls for staining specificity. Anti-Granzyme B (clone D2H2F) Rabbit mAb (Cell Signaling Cat. # 17215S) and rabbit anti-Ki67 antibody (Abcam, Cat# ab833) were used at 1:200 dilution. Goat anti-rabbit AF488 (Life Technologies, Cat. # A-11008) and Texas Red-X (ThermoFisher Cat. # T6391) secondary antibodies were used at 1:500 dilution in blocking buffer.

### Image acquisition and analysis

Tile-scan imaging of whole tumor sections (20×) was acquired in an Eclipse Ni-E Fluorescence Microscope (Nikon) with an ORCA-flash4.0 LT plus (Model C11440) Hamamatsu digital camera and an S Plan Fluor ELWD Ph1 ADM 20x objective (N/A 0.45 and refractive index of 1). Granzyme B imaging was performed in an LSM 800 Confocal Microscope (Carl Zeiss) using a Plan-Apochromat LCI Plan-Neofluar 25x (0.8 Imm Korr Water DIC) objective and a 63x/1.40 Oil DIC objective (Zeiss). For the images acquired with the Nikon Fluorescence microscope, the noise component was removed with Denoise AI utility, and the fluorescence background was subtracted with the NIS-Elements Advanced Research Analysis software (ver. 5.21.02, Nikon). Separated tile images per channel were converted to binary images, and automated image analysis was performed using Cell Profiler Software (ver. 4.0.7, Broad Institute MIT) [29], using a custom pipeline to determine the percent of nuclei (cells) with positive immunostaining for the marker over the total number of cells normalized to the area of the field of view, and expressed as per cent (%) of cells with positive marker/μm^2^. Immunofluorescence (IF) quantification data were obtained from individual tumors of at least three different mice (N=3) per genotype were at least two serial cryosections per tumor were imaged.

### Statistical Analysis

All the statistical analyses were performed using GraphPad Prism® ver. 8 (San Diego, CA, USA). Unless otherwise indicated, the results are expressed as the mean ± standard deviation (SD), and the biological (N) and technical replicates (n) for a given experiment are indicated in each figure. Animal numbers were estimated by *a priori* power analysis (G*Power ver.3.1 [30]) based on a pilot study of spontaneous tumor incidence between BPC-*Panx1*^+/-^ (heterozygous) and BPC-*Panx1*^−/-^ mice with an effect size of 1.0 and a power of 0.8. For multiple comparisons involving the genotype and sex of mice, two-way ANOVA was performed, followed by a Tukey-Kramer *post hoc* test for unequal sample sizes. For real-time qPCR, log_2_ (Relative Normalized expression) was used as data for statistical comparisons. Outliers in the gene expression data were detected using the ROUT method with a False Discovery Rate of less than 1% and excluded from analysis. Data normality was verified with the D’Agostino-Pearson test. One-way ANOVA followed by Sidak’s *post hoc* test was done to analyze relevant mean comparisons in the mRNA expression between tissues. For comparisons of IF quantification data (% cells with positive marker/μm^2^) per mouse were transformed using a Box-Cox transformation [31] and the means of the two groups (genotypes) were compared using a nested two-way ANOVA (Nested T-test). Statistical significance was considered when p < 0.05.

## RESULTS

### Germline *Panx1*-deletion does not reduce melanoma progression of Braf(V600E)/Pten(del) mice

To explore the *in vivo* effects of the *Panx1* global deletion on melanoma progression, we crossed *BRaf^CA/+^, Pten^LoxP/Loxp^, Tyr::CreER^T2^* (BPC) mice with the global *Panx1* knockout (*Panx1*^−/-^, *Panx1*KO) mouse line, reported on [27, 20]. In the BPC model, a 4-hydroxytamoxifen (4-HT)-inducible Cre recombinase is used for the melanocyte-specific expression of mutant BRaf^V600E^— a constitutively active protein serine kinase—and inactivation of the Pten phosphatase to produce primary cutaneous malignant melanomas (11). However, as in (23), we observed that spontaneous melanomas occurred in approximately 70% of the BPC colony in the absence of tamoxifen exposure. Thus, we started the 4-HT induction experiments in mice between 3–4 weeks of age that looked healthy and had no spontaneous melanoma formation.

Our results showed that regardless of the *Panx1* genotype, melanocytic pigmented lesions arose from the skin and quickly grew to form cutaneous melanomas as early as three weeks after topical application of 4-HT. Furthermore, although we applied 4-HT to a small zone in the dorsal skin of the lower back, we found that, in both animal groups, multiple tumor lesions grew not only in the application region but also in distant sites (Fig. 1A). As a result, all the visible melanoma lesions found in the dorsal and ventral/lateral skin of the animals were considered to determine tumor burden (Fig. 1B) and endpoint conditions. Overall, we observed that BPC-*Panx1*^−/-^ mice had similar tumor incidence (Fig. 1B), survival (Fig.1C), and mouse weight among sexes (Fig.1D) compared to *Panx1*-wildtype counterparts. However, although the total combined tumor growth over time (Fig.1E) was similar between genotypes, analysis of estimated tumor growth parameters showed sex differences of the tumor growth in BPC-*Panx1*^+/+^ mice (Fig. 1F and G). BPC-*Panx1*-wildtype male mice had a slightly lower (though not significant) tumor growth rate (Fig.1G, two-way ANOVA, Sex factor: F_1,23_ = 4.64, *p*<.05) with a significant increase (*p* = 0.188) in tumor doubling time (Fig.1F, two-way ANOVA, Sex factor: F_1,23_ = 7.86, *p*<.05) compared to female counterparts. However, this sex-driven difference was not significant in BPC-*Panx1*-deficient mice and overall tumor growth did not differ between genotypes. These results indicated that *Panx1* deletion in this mouse model did not reduce BRAF^V600E^/Pten-driven melanoma development.

**Figure 1.**
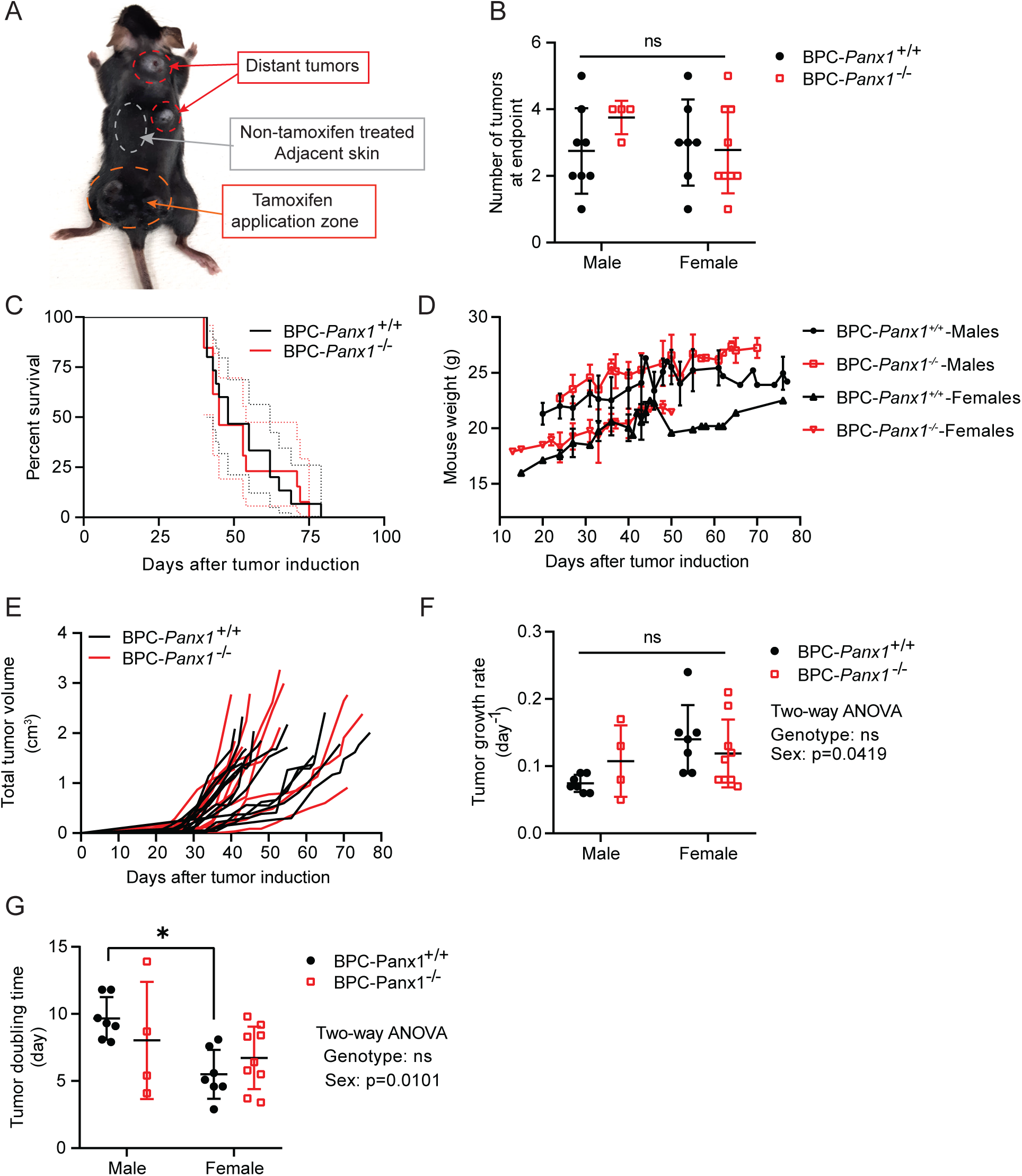
Global *Panx1* deletion does not improve survival or reduce tumor progression in the Braf(V600E)/Pten-deficient mouse melanoma model. **(A)** Representative picture of *Tyr::CreER; BRaf*^CA*/+*^*; Pten*^−/-^ (BPC) mice knockout (-/-) for *Panx1* with multiples tumors at endpoint. Region of the tamoxifen application is indicated in the picture. **(B)** Tumor incidence, recorded as the number of visible cutaneous melanoma lesions at endpoint, was similar between both mouse sexes and genotypes. **(C)** Kaplan-Meier survival curves were similar among BPC-*Panx1* wildtype (^+/+^) or BPC-*Panx1*^−/-^ mice (Log-rank test, p = 0.7803). Dotted lines denote 95% confidence intervals. **(D)** Comparison of mice body weight after tumor induction showed no differences among genotypes within the same sex group. Statistical significance was determined by multiple t-tests per timepoint using the Holm-Sidak method without assuming a consistent *SD*. **(E)** Tumor growth curves showed overlap between genotypes, with a group of mice exhibiting slightly delayed tumor growth. Individual lines depict measurements per mouse. **(F, G)** Tumor growth rate and doubling time were similar among genotypes but significantly different between males and females within the BPC-*Panx1*^+/+^ group. Two-way ANOVA showed no interaction between genotype and sex, but significant *p*-values for the sex factor in tumor doubling time (F_1,23_ = 7.855, p<.05) and growth rate (F_1,23_ = 4.642, p<.05), respectively. Symbols stand for individual data per mouse (BPC-*Panx1*^+/+^, n=15, and BPC-*Panx1*^−/−^, n=13) except for the graph in **D**, where mean ± *SD* are shown. Due to unequal sample sizes per sex, the Tukey-Kramer test was used as *post hoc* test for pairwise comparisons shown in **F** and **G**. Statistical significance was considered when *p*<0.05.

In general, most melanoma lesions were presented with a combination of melanotic (black) and amelanotic (white) regions (Fig. 2A). However, a few exceptions (less than 30% of animals per genotype) also had completely amelanotic tumors primarily distant from the tamoxifen application site. A classification of the tumors with the presence or complete absence of black pigmentation rendered no significant differences in their number among genotypes (Fig. 2A). All mice, regardless of the genotype, developed fast-growing tumors that, in most cases, were ulcerated by the time of endpoint (Fig. 2B). Since our previous reports indicated that PANX1 is highly expressed in human melanoma tumors compared to normal skin, we performed real-time-qPCR in matched samples of non-tamoxifen-treated (adjacent) skin and tamoxifen-induced tumors. *Panx1* mRNA expression was found to be 3.2-fold increased (Fig. 2C) in tumors of BPC-*Panx1*^+/+^ mice compared to skin, while undetectable in any tissue sample from BPC-*Panx1*^−/−^ mice, as expected. Moreover, mRNA expression of the *S100a1* melanoma marker [32, 33] had a 5.1-fold increase (p<0.05) in tumors of BPC-*Panx1*^−/−^ compared to the skin (Fig. 2D), indicating that the development of primary tumors occurred independently of *Panx1* expression.

**Figure 2.**
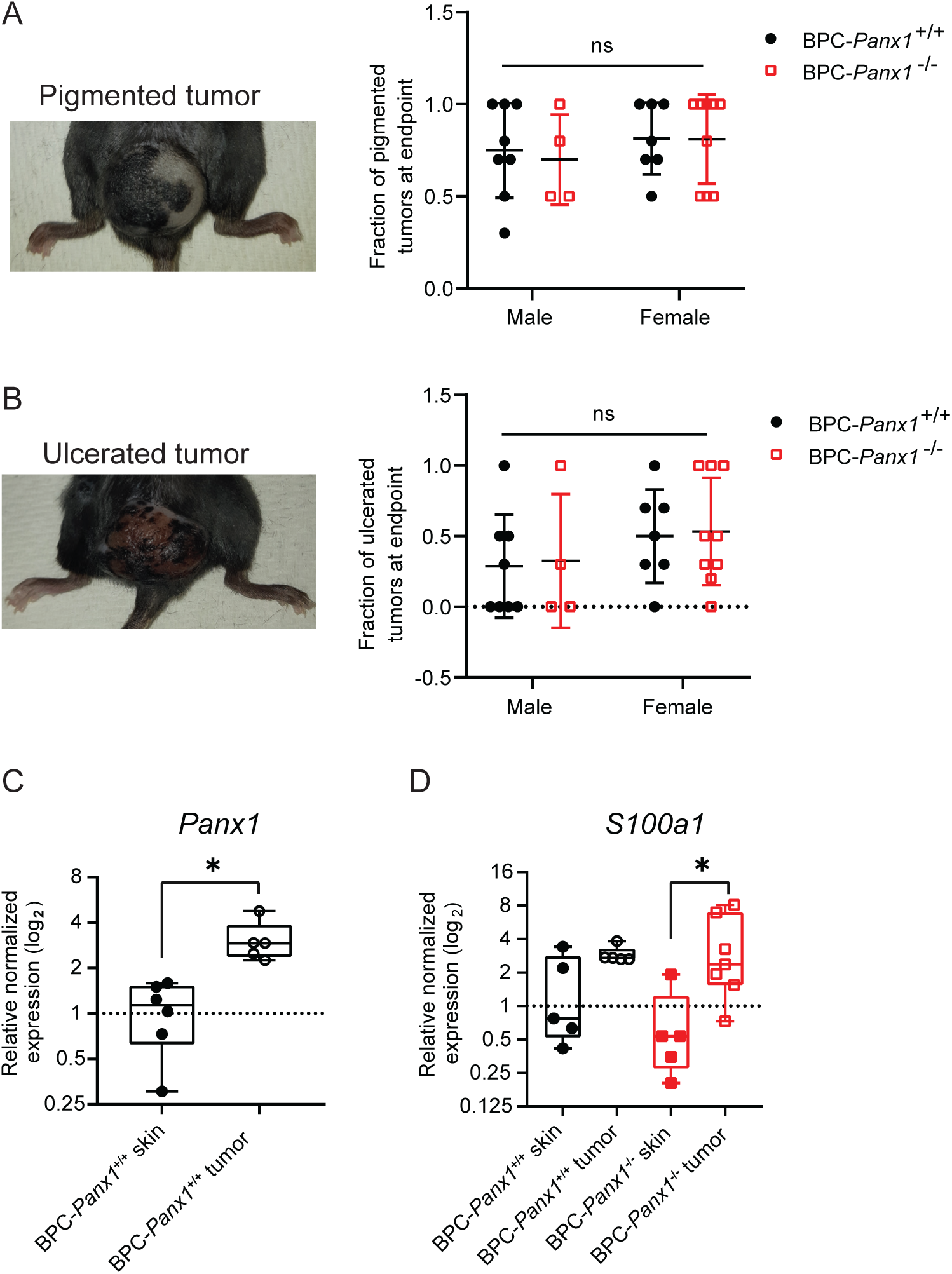
Pigmentation or ulceration of Braf(V600E)/Pten(del) melanoma lesions are not influenced by the global *Panx1* deletion. **(A, B)** Representative pictures of pigmented and ulcerated tumors. Fractions (out of the total tumors per mice) of pigmented or ulcerated tumors were similar among the genotype and sex groups. Symbols show individual data per mouse and horizontal lines, and error bars represent mean ± *SD*. Data were analyzed with two-way ANOVA. **(C, D)** mRNA expression of *Panx1* and melanoma marker *s100a1* in tumors compared to matched samples of non-tamoxifen treated skin per mouse. *Gapdh* and *18s* were used as reference genes, and the normalized expression is shown relative to one of the BPC-*Panx1*^+/+^ skin samples. Box plots represent the 95% confidence interval (CI) and the median (inner line), with whiskers representing the maximum and minimum values of the gene expression. Symbols display the individual expression data per mouse. Paired t-test was used to compare the log_2_ (normalized expression) of *Panx1* and one-way ANOVA followed by a Tukey test used to compare the expression of *S100a1*. At least N=5 paired samples were assayed by triplicate (n=3) for real-time-qPCR analysis. Statistical significance is shown as p<0.05(*).

### Braf(V600E)/Pten-driven melanoma metastasis to inguinal lymph nodes is not restricted by *Panx1*-deletion

Melanoma tumor cells are known for passing through the lymphatic system and entering the bloodstream to metastasize to other organs [34]. We observed localized and distant clusters of small black pigmented spots spread in the underside of dorsal skin that was not treated with tamoxifen, suggesting the presence of melanoma microlesions or that in-transit metastasis may have occurred (Fig. 3A). Regional black melanoma metastases were found in both right and left inguinal lymph nodes in both *Panx1* genotypes (Fig. 3B, C, and D) but this was not noticeable in major organs (e.g., lungs, liver, or brain).

**Figure 3.**
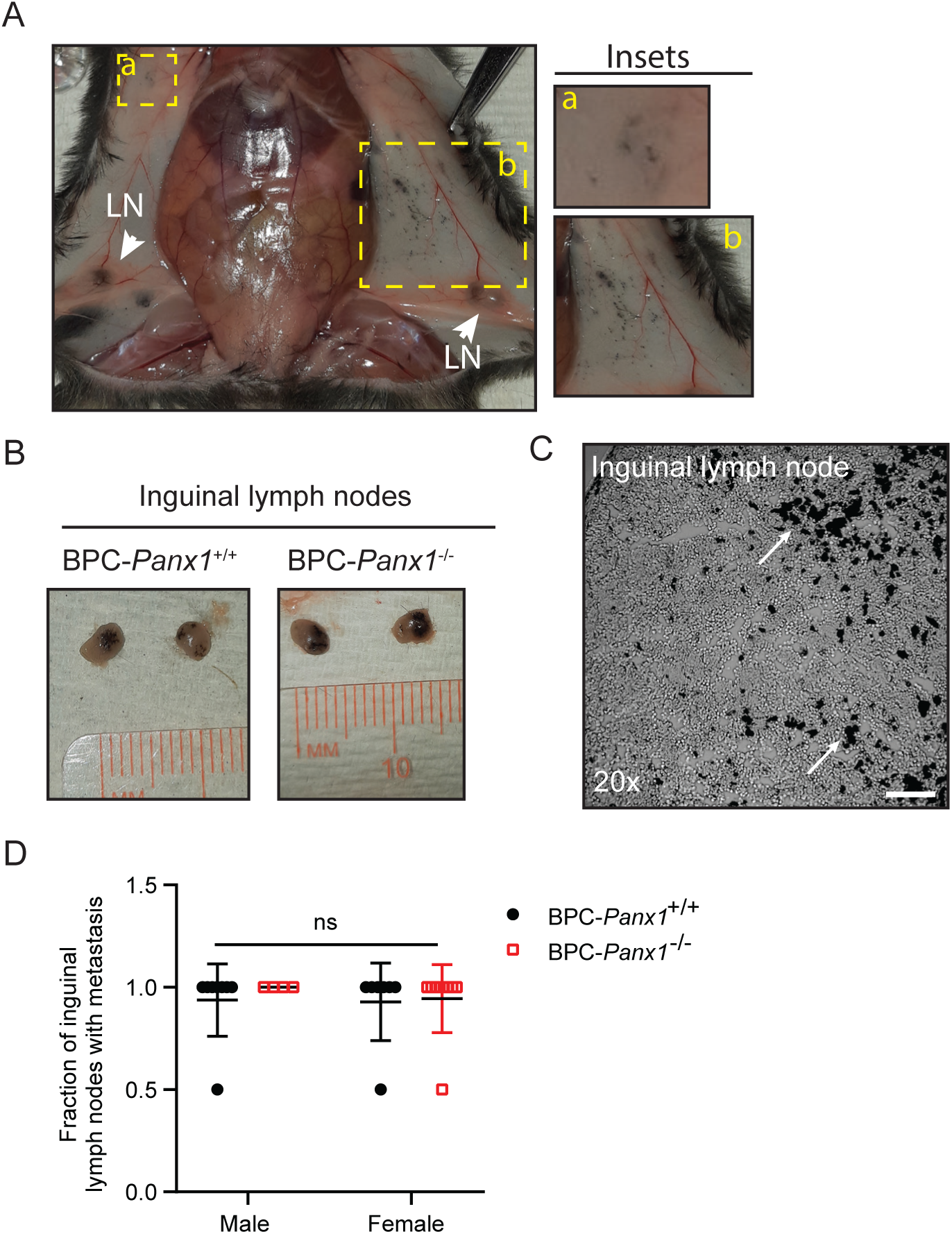
*Panx1* deletion does not prevent the development of Braf/Pten-driven melanoma micro-lesions or metastasis to inguinal lymph nodes in BPC mice. (**A**) Representative pictures of melanoma micro-lesions found underside the dorsal and ventral/lateral skin and located distant (inset **a**) and nearby (inset **b**) to the application sites of tamoxifen (e.g., lower back dorsal skin). LN: Inguinal draining lymph nodes. **(B)** Representative pictures of right and left inguinal lymph nodes showing black pigmentation indicative of melanoma metastasis. **(C)** Phase-contrast micrograph of an inguinal draining lymph node cryosection exhibiting melanin deposits and pigmented cells (white arrows). **(D)** Quantification of the fraction (out of two inguinal LN per mice) of draining lymph nodes classified with melanoma metastasis based on the presence or absence of visible pigmented lesions. Symbols correspond to the individual data per mouse where mean ± *SD* are shown. Data were analyzed with a two-way ANOVA. Statistical significance was considered when p<0.05.

### Spleens are significantly enlarged in females of tumor-bearing BPC-*Panx1*^−/-^ mice

Upon measuring the length and weight of spleens (Fig. 4A, B, C), a statistically significant sex-dependent difference was found in BPC-*Panx1*^−/-^ mice (two-way ANOVA, F_1, 20_ = 10.91, p<0.01). Females had an increased spleen weight index (normalized to body weight) compared to males of the BPC-*Panx1*^−/-^ cohort but this was not observed in BPC-*Panx1*^+/+^ mice or among genotypes. A positive correlation between spleen weight and length measurements (Fig. 4D) (Pearson’s coefficients r ≥ 0.7, *p*=0.0247 and *p*=0.0087) in both BPC-*Panx1*^+/+^ and BPC-*Panx1*^−/-^ mice, respectively, confirmed a similar trend in both parameters.

**Figure 4.**
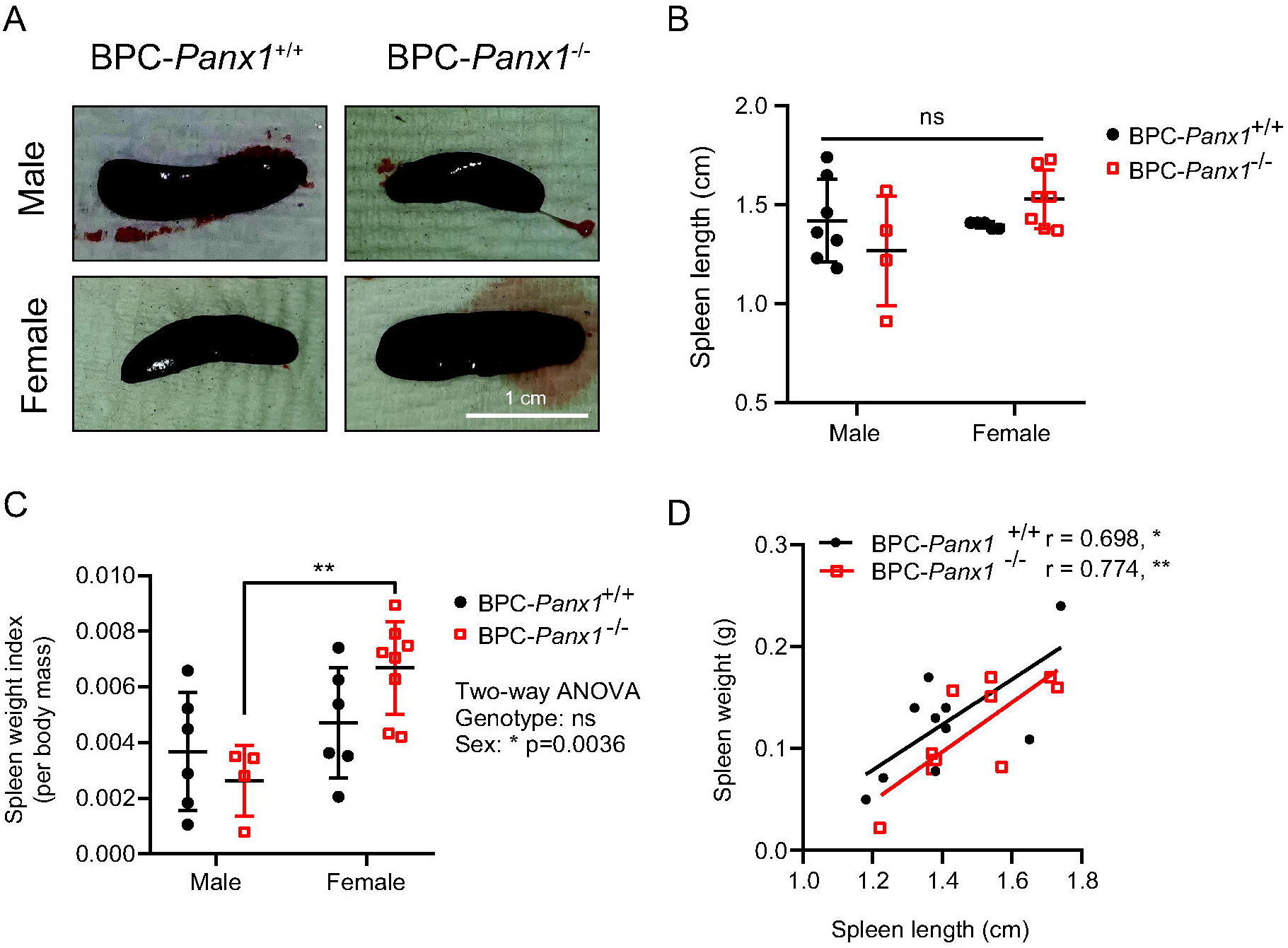
Sex-dependent differences in spleen size were observed for BPC-*Panx1*^−/−^ but not for BPC-*Panx1*^+/+^ mice. **(A)** Representative pictures of BPC mice spleens. **(B)** Morphological comparison of the spleen measured by length and **(C)** weight index (spleen weight normalized to body mass) showed a trend of enlarged spleens in female BPC-*Panx1*^−/-^ mice. Sex-dependent differences in BPC-*Panx1*^−/-^ mice were statistically significant (Two-way ANOVA, F_1, 20_ = 10.91, p<0.01). A Tukey-Kramer test used as *post hoc* test for pairwise comparisons showed a significantly (**, p<0.01) higher spleen weight index in BPC-*Panx1*^−/-^ females than males, but not in the BPC-*Panx1*^+/+^ cohort. Symbols correspond to data from individual mice where mean ± SD are shown. **(D)** A positive correlation was found between the spleen weight and length, confirming the trends of increased spleen size in female BPC-*Panx1*^−/-^ mice observed by both measurements.

### CD8 mRNA is significantly increased in *Panx1*-deficient skin and melanoma tumors

Due to PANX1′s role in the inflammatory response [17–22], we sought to examine changes in the infiltration of immune cells within the TME. We performed a real-time qPCR analysis of gene expression for markers of immune cell populations known to modulate the immune response against melanoma. We included primers specific to protein tyrosine phosphatase, receptor type, C (*Ptprc*) (also known as cluster of differentiation (CD) 45) for leukocytes; CD3 antigen, epsilon polypeptide (*CD3*ε) for T-lymphocytes; CD4 antigen (*CD4*) for T-helper lymphocytes; CD8 antigen, beta polypeptide (*CD8b*) for cytotoxic T cells; forkhead box P3 (*Foxp3*) for regulatory T cells (T-reg); natural cytotoxicity triggering receptor-1 or NCTR1 (*Ncr1*) of Natural killer (NK) cells and the cell surface glycoprotein F4/80 (*Agre1*) for monocytes/macrophages. The “immune” markers expression was evaluated in melanoma tumors and matched samples of non-tamoxifen-treated dorsal skin devoid of any visible melanoma lesions (referred to as “adjacent skin”). These adjacent skin samples were used to account for basal immune infiltration levels or skin-resident immune cells [35, 36] at the time of sample collection.

When analyzing tumors compared with their matched skin samples, we observed a significant (F_3, 20_ = 6.24, p<0.01) 4.3 fold-increase expression of CD45 only within the BPC-*Panx1*^+/+^ mice group (Fig. 5A). The F4/80 mRNA expression (F_3, 18_ = 19.38, p<0.0001) was 14.0 (p<0.0001) and 7.7-fold (p<0.01) higher in tumors than the skin in BPC-*Panx1*^+/+^ and BPC-*Panx1*^−/-^ mice, respectively (Fig. 5B). On the other hand, transcripts of NCTR1 (F_3, 11_ = 8.24, p<0.01) (Fig. 5C) were 9.3-fold upregulated (p<0.01) in BPC-*Panx1*^−/−^ tumors versus adjacent skin. *CD3*ε (Fig. 5D) showed a similar expression regardless of the sample type or *Panx1* genotype. While *CD4* mRNA expression (F_3, 11_ = 27.46, p<0.0001) was significantly augmented in the tumors compared to the skin (Fig. 5E) with a 10.5 (p<0.01) and 18.9-fold (p<0.001) increase in the BPC-*Panx1*^−/-^ and BPC-*Panx1*^+/+^, respectively; and 5-fold upregulated (p<0.05) in the skin of BPC-*Panx1*^−/-^ vs. that of BPC-*Panx1*^+/+^. Moreover, *Foxp3* (F_3, 16_ = 9.80, p<0.001) was upregulated (5.9-fold, p<0.05, and 15.2-fold, p<0.01) in tumors of BPC-*Panx1*^+/+^ and BPC-*Panx1*^−/-^ mice, respectively (Fig 5G). Notably, *CD8* (F_3, 16_ = 15.39, p<0.0001) was significantly increased (p<0.01) in the skin and tumors (9- and 9.5-fold, respectively) of BPC-*Panx1*^−/-^ compared to BPC-*Panx1*^+/+^ animals (Fig. 5F). Taken together, these results indicated that *Panx1* deficiency did not impair the overall recruitment of immune cells to the primary tumors in this melanoma model but had an effect on increasing the transcript expression of CD8 in skin and tumors of BPC-mice.

**Figure 5.**
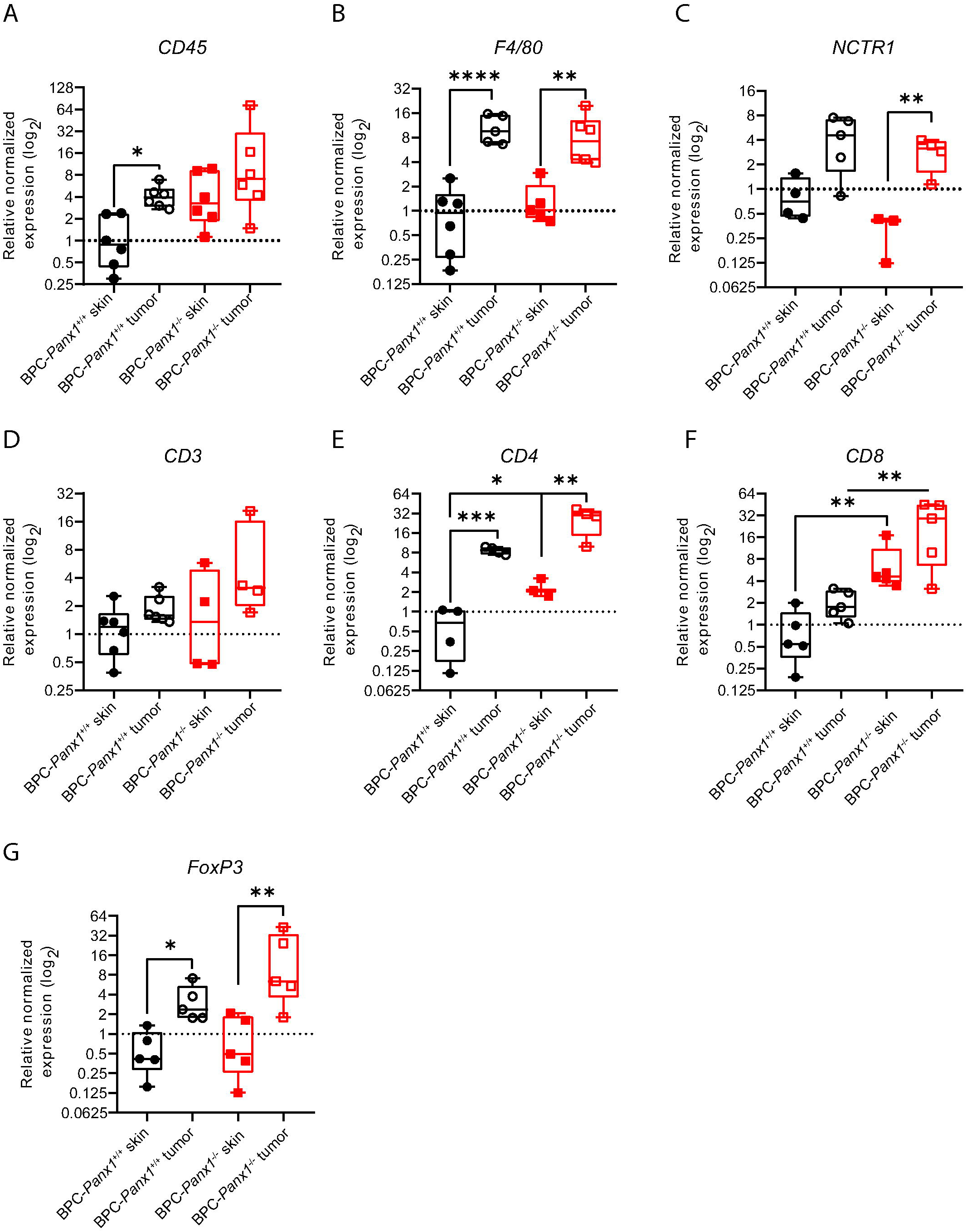
CD8 T-lymphocyte marker is significantly increased in the non-tamoxifen treated skin and melanoma tumors of BPC-mice lacking PANX1. **(A-G)** Real-time qPCR analysis of immune cell markers expression in tamoxifen-induced tumors and matched non-tamoxifen treated skin. *Gapdh* and *18s* were used as reference genes, and the normalized expression was calculated relative to one of the BPC-*Panx1*^+/+^ skin samples. Box plots represent the 95% confidence interval (CI) and the median (inner line), with whiskers showing the maximum and minimum values of the gene expression. Each sample was assayed by triplicate (n=3). The symbols represent the expression data of each mouse (at least N=3 per genotype). The log_2_ of the expression per group was analyzed by one-way ANOVA followed by a Sidak *post hoc* test. Only significant *p* values of relevant comparisons between groups are shown as *p*<0.05 (*), <0.01(**), <0.001(***).

Given the similarity in tumor burden among genotypes but increased CD8 mRNA in the BPC-*Panx1*^−/-^ tumors, we investigated if there were differences in the activation/exhaustion phenotype of TILs and immunosuppression compared to those of the BPC-*Panx1*^+/+^ cohort. We observed a trend of lower transcript expression (not significant) of *CD69* (early T cell activation marker, [37]) in BPC-*Panx1*^−/-^ tumors (Fig. 6A). Moreover, no differences in expression of the regulatory immune checkpoint receptor lymphocyte activation gene-3 (*LAG-3*) [38] or programmed cell death protein-ligand 1(PD-L1) [39] were found between genotypes (Fig.6B, C).

**Figure 6.**
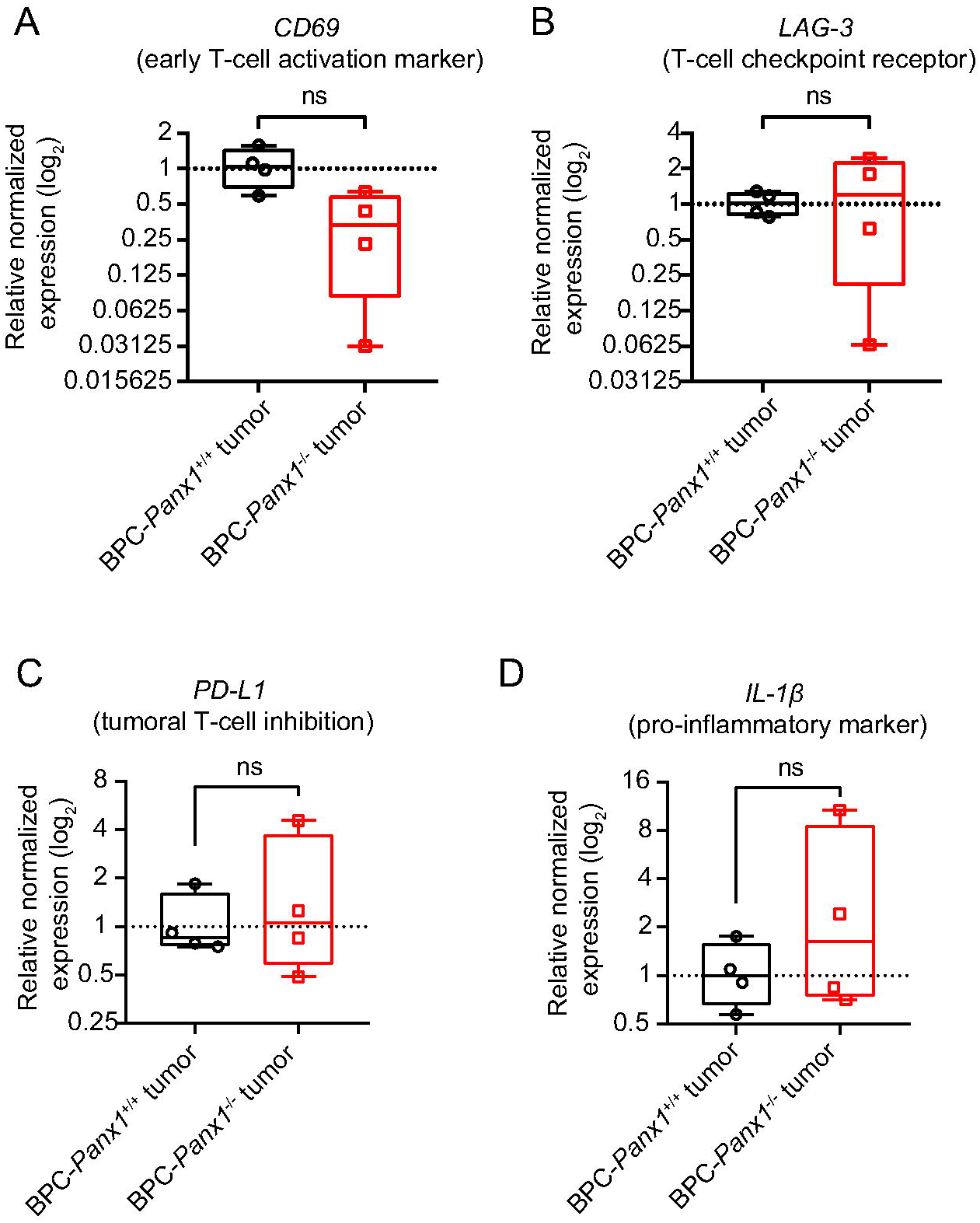
Global *Panx1-*deletion does not affect the activation phenotype of tumor-infiltrated T-lymphocytes. Real-time qPCR analysis of **(A)** the early T cell immune activation marker *CD69*, **(B)** checkpoint receptor lymphocyte activation gene-3 (*LAG-3*), **(C)** immune checkpoint molecule programmed cell death 1-ligand (*PD-L1*), and **(D)** *IL-1*β in tumor samples. *Gapdh* was used as a reference gene, and the normalized expression is relative to one of the BPC-*Panx1*^+/+^ skin samples. Box plots represent the 95% confidence interval (CI) and the median (inner line), with whiskers showing the maximum and minimum values of the gene expression. Each sample was assayed by triplicate (n=3), and the symbols represent the expression data of each mouse (at least N=4 per genotype). A t-test compared the mean of the log_2_ of the relative expression per group; statistical significance was considered when *p*<0.05 (*).

Finally, due to the association of PANX1 channel function and the pro-inflammatory and tumor promoter cytokine: interleukin-1β (IL-1β), we investigated whether the *IL-1*β expression was affected within the TME of BPC *Panx1*-deficient mice. IL-1β mRNA levels (Fig. 6D) were found to be highly variable in the BPC-*Panx1*^−/-^ tumors but were not different compared to tumors of the BPC-*Panx1*^+/+^ mice.

### *Panx1*-deletion in tumors does not impair infiltration of CD4^+^, CD8^+^ T lymphocytes, and Granzyme-B producing cells

To confirm the differences found in mRNA markers for T cell infiltration in tumors between *Panx1* genotypes we performed IF staining on PLP-fixed tumor cryosections to verify presence of CD4^+^, CD8^+^, and Foxp3^+^ lymphocytic cells (Fig. 7A). Overall, CD4^+^ and CD8^+^ T cells were seen in small but sparsely distributed clusters closer to the skin epidermis of the primary melanoma tumors, and Foxp3^+^ T cells showed a more homogeneous distribution within the tumor core irrespective of genotype (Fig. 7A). Overall, the quantification of the immune cell infiltration of lymphocytes agreed with the previous results on real-time qPCR immune markers. The percent of positive CD4^+^ and CD8^+^ T cells was variable among tumor samples and, although not statistically significant, it trended increased in BPC-*Panx1*^−/-^ vs BPC-*Panx1*^+/+^ tumors with a median value of 3.907 (interquartile range (IQR) 0.0-11.80) vs 1.523 (IQR 0.0-4.947) % CD4^+^cells (x 10^6^) /μm and 4.163 (IQR 0.0-4.163) vs 0.3672 (IQR 0.0-0.367) % CD8^+^cells (x 10^6^) /μm, respectively (Fig.7B,C). Moreover, the median percent of Foxp3^+^ T cells in was similar among genotypes with 6.046 (IQR 0.538-1.328) vs 1.477 (IQR 0.0-12.06) (x 10^6^) /μm in BPC-*Panx1*^+/+^ vs BPC-*Panx1*^−/-^ tumors (Fig.7D) . Lastly, we assessed the IF staining of Granzyme B (GzmB) to determine whether differences in cytotoxicity of T- or NK cells could be detected within the TME. A trend (not statistically significant, Nested T-test p>0.05) of increased of % GzmB^+^ cells (x 10^6^) /μm was observed in tumors of BPC-*Panx1*^−/-^ mice with median value of 14.30 (IQR 5.570-60.20) compared to 3.250 (IQR 0.0-7.280) in BPC-*Panx1*^+/+^ mice, with GzmB detection mostly localized as intracellular granules (Fig. 7E, F). These results correlate with the infiltration of CD8^+^ T cells and, likely, NK cells since GzmB can be abundant as cytosolic granules in both cytolytic immune cell types.

**Figure 7.**
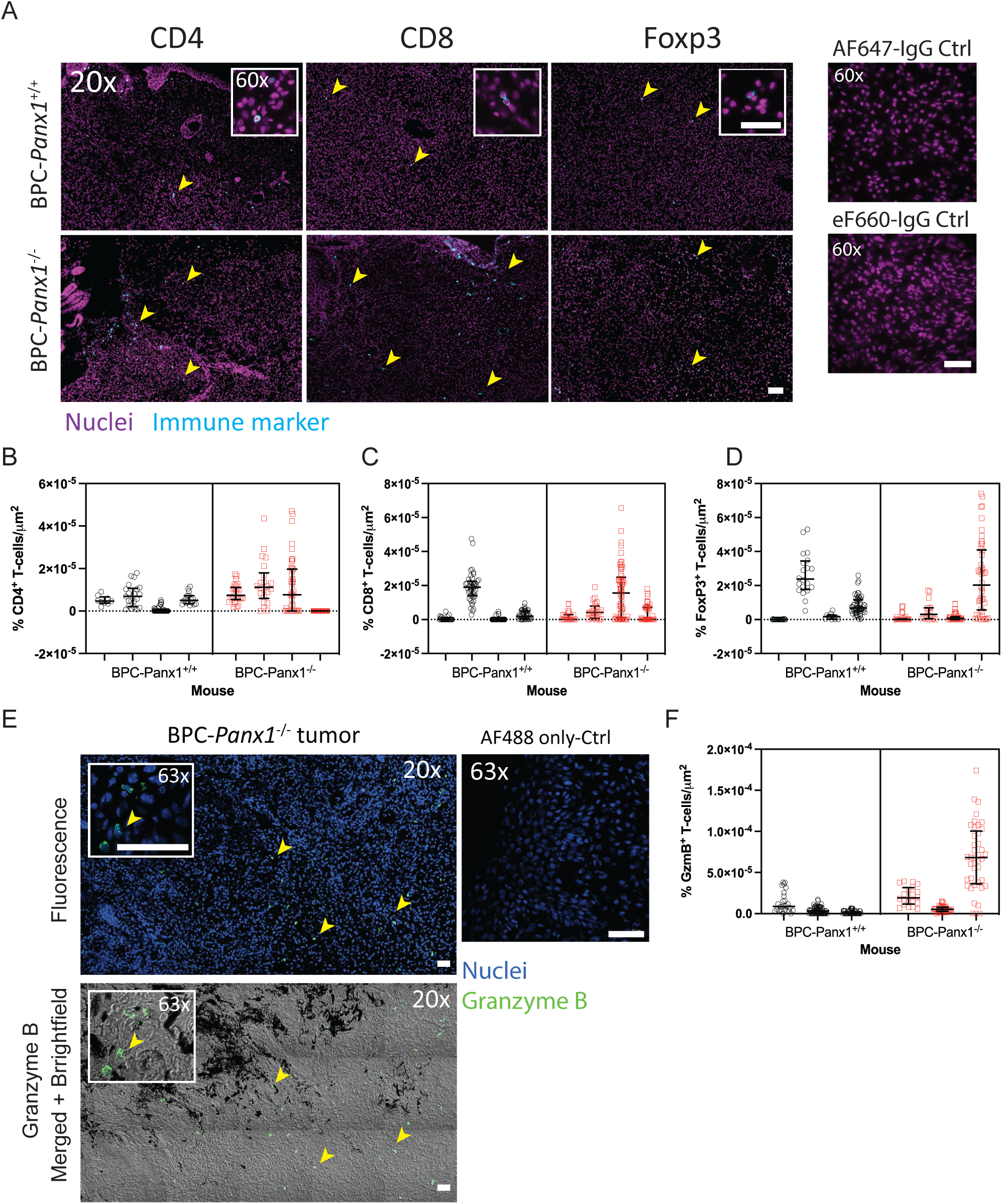
Tumor-infiltrating CD4^+^, CD8^+^ T-lymphocytes, and Granzyme B producing cells are present in BPC-*Panx1*^−/-^ tumors. **(A)** Representative micrographs (20x, insets at 60x magnification) showing immunofluorescence staining of primary melanoma tumor cryosections with immune cell infiltration markers for CD4+, CD8+, and FoxP3+ T-lymphocytes (shown in cyan). Nuclei (in magenta) was counterstained with Hoechst 33324. **(B-D)** Immunofluorescence quantification of CD4, CD8, and FoxP3 positive cells for the whole tumor section per animals **(E)** Representative granzyme B (GzmB) immunofluorescence staining of a BPC-*Panx1*^−/-^ tumor (fluorescence channels on top and brightfield channel at the bottom) with the intra-tumoral distribution of GzmB+ cells and black pigmented regions in the tumor. Nuclei are shown in blue, and GzmB is shown in green. Yellow arrowheads indicate stained immune cells. Scale bars = 40 µm. **(F)** Quantification of GzmB immunostaining. The percent of nuclei with positive immune marker staining was normalized per area of the field of view (FOV). In the nested graphs, at least three mice (N=3) per genotype are shown. Lines within graphs depict the median with the interquartile range, and individual measurements per FOV are shown with symbols. Box-Cox transformed data were compared using a two-tailed Nested T test. Statistical significance was considered when *p*<0.05.

Collectively, these results confirmed that the deletion of *Panx1* did not affect the infiltration of CD4^+^ and CD8^+^ T lymphocytes in the tumors, although the cytotoxic effect seemed to be insufficient to prevent the tumor progression in BPC-*Panx1*^−/-^ mice.

## DISCUSSION

PANX1 channels have attracted considerable attention due to their multiple roles in inflammation, cell death, and cancer [18, 40]. We have demonstrated that PANX1 influences the regulation of effector molecules that control melanocyte differentiation (e.g., MITF and β-catenin) and the metabolism of melanoma cells [15, 14, 16]. Furthermore, we have shown promising results after silencing *Panx1* expression and reducing the growth and tumorigenicity of melanoma cell lines *in vitro*. In this work, using the immunocompetent BPC mouse melanoma model, we did not observe a reduction in Braf(V600E)/Pten(del)-driven melanomas or improve mice survival upon germline *Panx1* deletion. Of note, our previous *in vitro* and *ex vivo* work targeting PANX1 in melanoma employed murine B16-F0, -F1, -F10 and human BRAF(V600E)-mutant melanoma cell lines (A375-P and A375-M2) that do not entirely share the same genomic alterations of the BPC model [15, 41]. The deletion of *Pten* is known to have a synergistic effect and dramatically accelerates BRAFV600E-induced melanoma tumors in BPC mice [26, 27, 42]. Therefore, we hypothesize that oncogenic mutations intrinsic to this mouse BPC model may have overridden our anticipated anti-tumoral effect of *Panx1*-deletion. Since this is a highly penetrant mouse model, a syngeneic melanoma model may be more suitable to study the effect of *Panx1*-deletion in melanoma metastasis [43]. Besides, the germline deletion of *Panx1* does not necessarily imply the same effect previously observed with the use of PANX1 channel blockers (e.g., with carbenoxolone, probenecid, or spironolactone) due to possible compensation with other proteins (e.g., connexins or other paralogs) yet to be tested using this or another *in vivo* melanoma model. We also observed the formation of micro-metastasis to draining inguinal lymph nodes (a phenotype reported elsewhere [26, 27, 44, 28]) in *Panx1*-deficient mice. Therefore, we concluded that *Panx1*-deletion does not control the early spread of melanoma. Due to the ubiquitous expression of *Panx1*, its global deletion should have affected many cell types, including melanoma and immune cells which may resemble the effect of an irreversible depletion of PANX1 function in which adaptative or alternative molecular mechanisms are in play that helped to sustain tumor growth.

Interestingly, we noticed a sex dimorphism in melanoma tumor progression where BPC-*Panx1*^+/+^ female mice seemed to have a more aggressive disease (Fig.1E, G) than males of the same cohort. Other researchers have noted this outcome in a C57B/6J background, but the origin of this difference has not been clarified yet [45]. It has been speculated that this may be due to the presence of alternative steroid receptors in melanoma cells or technical limitations with tamoxifen (an estrogen receptor modulator) differentially influencing the mouse sexes in this melanoma model [45]. This sex-dependency was lost in BPC-*Panx1*^−/-^ mice, suggesting that *Panx1*-deletion may influence such a sex-driven phenotype. It is known that sexual dimorphism exists in the response of *Panx1*-deficient mice in ischemia [46] and epilepsy [47], and research on this is still lacking in the melanoma and immune cell recruitment contexts. A recent study has shown that tumor progression and anti-tumoral T cell response are sex-dependent in a syngeneic B16-F10/BL6 mouse model, however, in contrast to our observation, such study showed that females had less tumor growth rate and higher CD4^+^ and CD8^+^ T cell infiltrates compared to males [48]. Due to the lack of sample availability at the time of the experiment, we did not quantify differences in T cell subtypes between samples from different sexes. Therefore, it is unclear the nature of the loss of such sex disparity and further investigation may be required to clarify this finding.

To date, there are no reports of major abnormalities or immune-related diseases in the Panx1-deficient mice (reviewed at [18]). Here, we report for the first time that tumor-bearing BPC-*Panx1*^−/-^ female mice have significantly augmented spleen size compared to their male counterparts. Due to the short timeframe for tumor growth (∼2 months) and given that the sexual dimorphism was found only in BPC-*Panx1*^−/-^ spleens, we speculate that characteristic developmental differences may account for this phenotype in *Panx1*^−/-^ mice. Nonetheless, further research is warranted to determine whether global *Panx1*-deletion alone exerts sex-specific changes in the spleen morphology and whether this causes an immune dysregulation that could impact the tumor response in mice.

Tumor-infiltrating lymphocytes (TILs) have already been described in this BPC model [28]. However, others have highlighted that this model does not appear to induce a marked immune response due to the lack of additional tumoral somatic mutations/neoepitopes [43]. Notably, our results showed trends of increased TILs in BPC-*Panx1*^−/-^ tumors which suggests that Panx1 has an influence on the immune infiltration in the TME of this model.

It must be taken into consideration that, although not investigated here, in the TME, there are diverse immunosuppressive cells like myeloid-derived suppressor cells, T regulatory (Treg) cells, tumor-associated fibroblasts and macrophages. These cells release various soluble factors, including reactive oxygen species (ROS), which effectively hinder the response of other effector cells like natural killer (NK) cells. Moreover, elevated levels of fibroblasts in the TME contribute to increased secretion of metalloproteinases, leading to the further shedding of ligands that could otherwise engage with NK cells [49]. Additionally, tumor-associated fibroblasts are known to negatively influence NK cells by inhibiting the upregulation of activating receptors induced by cytokines [50]. Furthermore, immunosuppression of NK cells can be exerted by exosomes derived from melanoma [51], which were also not considered in the present study.

Despite our limited analysis on immunosuppression, we found no difference in FoxP3 mRNA expression and T-reg cells infiltration between tumors from both mice cohorts but an increase of FoxP3 transcript expression in tumors (Figures 5G and 7D) was evident compared to the skin tissue. This suggest that deletion of *Panx1* alone may not influence T-reg-cell driven immunosuppression in this tumor model. On the other hand, myeloid-derived suppressor cells, along with high PD-1 expression on the T cells, are known to highly immunosuppress this melanoma model [28]. It is unclear if these or other subsets of CD4^+^ T cells (Th1 or Th17) and macrophages (M2), may have contributed to immune evasion. Furthermore, we found no differences in gene-expression of markers of TIL activation (CD69)[52, 53], exhaustion (LAG-3), or tumor immune evasion (PD-L1), however, our strategy could not provide a definitive assessment of the proportion of the activation/exhaustion status within the TIL subset. Nevertheless, it remains to be tested whether the absence of PANX1 in a specific subtype of the immune cell in the tumor could be exploited to counterbalance the immunosuppressive or evasive mechanisms of melanoma.

PANX1 has been proposed as a potential target to regulate antitumor immunity in melanoma due to the PANX1-mediated release of proinflammatory cytokine IL-1β (reviewed in [54]). IL-1β in melanoma elicits inflammation and the expansion of highly immunosuppressive myeloid-derived suppressor cells [55]. In this study, we found no significant differences in IL-1β mRNA expression among cohorts (Fig. 6D) suggesting that this immune evasion mechanism may be still in place contributing to tumor progression despite the increased TIL in Panx1^−/-^ tumors. In addition, using this same *Panx1*^−/-^ mouse strain, another group showed that marrow-derived macrophages have no difference in the secretion of IL-1β or inflammasome activation [20]. This underscores the need for research into the crosstalk between immune and tumor cells under cell-specific PANX1 suppression/inhibition to better understand the implications of PANX1 targeting in the context of melanoma and immunotherapy. As a limitation of this study, *Panx1* gene deletion was not directed to any specific cell compartment, therefore it is difficult to determine the specific impact of this deletion in the immune and tumor compartments distinctly. A possible solution to this problem could include the use of a syngeneic mouse model where Panx1 deletion can be targeted to specific immune cell types and be studied in a more relevant immunogenic melanoma model like YOVAL1.1 [56].

This transgenic melanoma mouse model was originally developed to harness two significant oncogenic alterations (BRAF^V600E^ and PTEN (loss)), but it is not representative of the larger patient population since in our analysis of the Cutaneous Melanoma TCCGA PanCancer data, we found that only 5.0% of cases had co-occurrence of both mutations (Supplemental Fig. S1A). Furthermore, since *PANX1* expression seems independent of genomic alterations of melanoma (Supplemental Fig. S1B) and is present at all stages of the disease [15], a larger patient population may benefit from PANX1 targeted therapy. Interestingly, *Pten*-null cancers seem resistant to immune checkpoint inhibitory antibodies, where PTEN loss in tumor cells increases the expression of immunosuppressive cytokines inhibiting trafficking and T cell-mediated killing of tumors [57] [58]. Hence, our results point towards the direction of targeting PANX1 as a potential alternative to overcome T cell exclusion in immunosuppressive TMEs.

In conclusion, our study demonstrates that a *Panx1* deletion cannot overcome the aggressive tumorigenic effect of Braf(V600E)/Pten(del) driver mutations, but it increases the tumor T cell infiltration with a preference in CD8^+^ cytotoxic T cells. This could be advantageous since a considerable therapeutic benefit is achieved in patients with a pre-existing CD8^+^ T cell infiltration within the melanoma microenvironment [59]. Future preclinical studies should include the combination strategies in which PANX1 inhibition could be considered as means to stimulate cytotoxic T-cell tumor infiltration to improve the efficacy of immunotherapies.

## Acknowledgments

The authors thank Dr. Saman Maleki-Vareki and Dr. Frank Beier for their input on the article. This study was supported by a Canadian Institutes of Health Research CIHR Project grant to Silvia Penuela and Lina Dagnino (FRN 153112)

## Authors’ Contributions

**S. Penuela:** Supervision, conceptualization, data analysis, funding acquisition, and writing; **RE Sanchez-Pupo:** Performed most experimental procedures, data analysis, visualization, methodology, wrote and edited the original draft; **G. Finch:** Data curation, spleen measurements, qPCR experiments, and manuscript editing; **D. Johnston:** Technical assistance, resources management, tumor monitoring, sample collection, colony breeding, and dissections; **R. Abdo:** Dissections and sample collection; **S. Kerfoot, H. Craig:** Collaboration, provided IF protocols, input for experiments and writing; **L. Dagnino:** Collaboration with grant funding and input on the writing and data analysis.

The authors declare no conflicts of interest of any kind with the current manuscript.

**Supplemental Figure S1. PANX1 mRNA expression is similar between patients with and without co-occurrence of BRAF(V600E) /Pten (homozygous deletion)**. **(A)** Analysis of Skin Cutaneous Melanoma TCGA PanCancer data shows a low proportion (∼5.0%) of patients have co-occurrence of the BRAF(V600E) and homozygous deletion of PTEN. **(B)** Both cohorts of patients exhibit similar levels (p=0.4970) of PANX1 mRNA expression. Similar **(C)** overall and **(D)** progression-free survival curves exist between patients with (red line; n = 144) or without (blue line; n = 19) co-occurrence of BRAF(V600E) mutation/Pten (deep deletion). Data in graphs were exported from cBioPortal (as of Jan/12/2021)(49, 50). RNASeqV2 data (Batch normalized from Illumina HiSeq) is derived from TCGA PanCancer Atlas datasets, processed, and normalized using RSEM. Bar and error graphs stand for the geometric mean with 95% CI, and individual patient expression data is depicted with symbols. A two-tailed unpaired t-test with Welch’s correction was used to compare the means of log_2_(mRNA PANX1 expression), and a log-rank test was used to determine statistical significance (p<0.05) in the Kaplan-Meyer curves.

## REFERENCES

1. Tas F (2012) Metastatic behavior in melanoma: timing, pattern, survival, and influencing factors. J Oncol 2012, 647684, doi: 10.1155/2012/647684.

2. Boussios S, Rassy E, Samartzis E, Moschetta M, Sheriff M, Pérez-Fidalgo JA & Pavlidis N (2021) Melanoma of unknown primary: New perspectives for an old story. Crit Rev Oncol Hematol 158, 103208, doi: 10.1016/j.critrevonc.2020.103208.

3. Carlino MS, Larkin J & Long GV (2021) Immune checkpoint inhibitors in melanoma. Lancet 398, 1002–1014, doi: 10.1016/S0140-6736(21)01206-X.

4. Stahl JM, Sharma A, Cheung M, Zimmerman M, Cheng JQ, Bosenberg MW, Kester M, Sandirasegarane L & Robertson GP (2004) Deregulated Akt3 activity promotes development of malignant melanoma. Cancer Res 64, 7002–7010, doi: 10.1158/0008-5472.CAN-04-1399.

5. Birck A, Ahrenkiel V, Zeuthen J, Hou-Jensen K & Guldberg P (2000) Mutation and allelic loss of the PTEN/MMAC1 gene in primary and metastatic melanoma biopsies. J Invest Dermatol 114, 277–280, doi: 10.1046/j.1523-1747.2000.00877.x.

6. Adeleke S, Okoli S, Augustine A, Galante J, Agnihotri A, Uccello M, Ghose A, Moschetta M & Boussios S (2023) Melanoma-the therapeutic considerations in the clinical practice. Ann Palliat Med, doi: 10.21037/apm-22-1432.

7. Dempke WCM, Fenchel K, Uciechowski P & Dale SP (2017) Second- and third-generation drugs for immuno-oncology treatment-The more the better? Eur J Cancer 74, 55–72, doi: 10.1016/j.ejca.2017.01.001.

8. Havel JJ, Chowell D & Chan TA (2019) The evolving landscape of biomarkers for checkpoint inhibitor immunotherapy. Nat Rev Cancer 19, 133–150, doi: 10.1038/s41568-019-0116-x.

9. Wolchok JD, Chiarion-Sileni V, Gonzalez R, Rutkowski P, Grob JJ, Cowey CL, Lao CD, Wagstaff J, Schadendorf D, Ferrucci PF, Smylie M, Dummer R, Hill A, Hogg D, Haanen J, Carlino MS, Bechter O, Maio M, Marquez-Rodas I, Guidoboni M, McArthur G, Lebbe C, Ascierto PA, Long GV, Cebon J, Sosman J, Postow MA, Callahan MK, Walker D, Rollin L, Bhore R, Hodi FS & Larkin J (2017) Overall Survival with Combined Nivolumab and Ipilimumab in Advanced Melanoma. N Engl J Med 377, 1345–1356, doi: 10.1056/NEJMoa1709684.

10. Jerby-Arnon L, Shah P, Cuoco MS, Rodman C, Su MJ, Melms JC, Leeson R, Kanodia A, Mei S, Lin JR, Wang S, Rabasha B, Liu D, Zhang G, Margolais C, Ashenberg O, Ott PA, Buchbinder EI, Haq R, Hodi FS, Boland GM, Sullivan RJ, Frederick DT, Miao B, Moll T, Flaherty KT, Herlyn M, Jenkins RW, Thummalapalli R, Kowalczyk MS, Canadas I, Schilling B, Cartwright ANR, Luoma AM, Malu S, Hwu P, Bernatchez C, Forget MA, Barbie DA, Shalek AK, Tirosh I, Sorger PK, Wucherpfennig K, Van Allen EM, Schadendorf D, Johnson BE, Rotem A, Rozenblatt-Rosen O, Garraway LA, Yoon CH, Izar B & Regev A (2018) A Cancer Cell Program Promotes T Cell Exclusion and Resistance to Checkpoint Blockade. Cell 175, 984–997 e924, doi: 10.1016/j.cell.2018.09.006.

11. Cabrita R, Mitra S, Sanna A, Ekedahl H, Lovgren K, Olsson H, Ingvar C, Isaksson K, Lauss M, Carneiro A & Jonsson G (2020) The Role of PTEN Loss in Immune Escape, Melanoma Prognosis and Therapy Response. Cancers (Basel) 12, doi: 10.3390/cancers12030742.

12. Trujillo JA, Luke JJ, Zha Y, Segal JP, Ritterhouse LL, Spranger S, Matijevich K & Gajewski TF (2019) Secondary resistance to immunotherapy associated with beta-catenin pathway activation or PTEN loss in metastatic melanoma. J Immunother Cancer 7, 295, doi: 10.1186/s40425-019-0780-0.

13. Medina CB, Mehrotra P, Arandjelovic S, Perry JSA, Guo Y, Morioka S, Barron B, Walk SF, Ghesquiere B, Krupnick AS, Lorenz U & Ravichandran KS (2020) Metabolites released from apoptotic cells act as tissue messengers. Nature 580, 130–135, doi: 10.1038/s41586-020-2121-3.

14. Penuela S, Gyenis L, Ablack A, Churko JM, Berger AC, Litchfield DW, Lewis JD & Laird DW (2012) Loss of pannexin 1 attenuates melanoma progression by reversion to a melanocytic phenotype. J Biol Chem 287, 29184–29193, doi: 10.1074/jbc.M112.377176.

15. Freeman TJ, Sayedyahossein S, Johnston D, Sanchez-Pupo RE, O’Donnell B, Huang K, Lakhani Z, Nouri-Nejad D, Barr KJ, Harland L, Latosinsky S, Grant A, Dagnino L & Penuela S (2019) Inhibition of Pannexin 1 Reduces the Tumorigenic Properties of Human Melanoma Cells. Cancers (Basel) 11, doi: 10.3390/cancers11010102.

16. Sayedyahossein S, Huang K, Li Z, Zhang C, Kozlov AM, Johnston D, Nouri-Nejad D, Dagnino L, Betts DH, Sacks DB & Penuela S (2021) Pannexin 1 binds beta-catenin to modulate melanoma cell growth and metabolism. J Biol Chem, 100478, doi: 10.1016/j.jbc.2021.100478.

17. Chekeni FB, Elliott MR, Sandilos JK, Walk SF, Kinchen JM, Lazarowski ER, Armstrong AJ, Penuela S, Laird DW, Salvesen GS, Isakson BE, Bayliss DA & Ravichandran KS (2010) Pannexin 1 channels mediate ‘find-me’ signal release and membrane permeability during apoptosis. Nature 467, 863–867, doi: 10.1038/nature09413.

18. Crespo Yanguas S, Willebrords J, Johnstone SR, Maes M, Decrock E, De Bock M, Leybaert L, Cogliati B & Vinken M (2017) Pannexin1 as mediator of inflammation and cell death. Biochim Biophys Acta Mol Cell Res 1864, 51–61, doi: 10.1016/j.bbamcr.2016.10.006.

19. Douanne T, Andre-Gregoire G, Trillet K, Thys A, Papin A, Feyeux M, Hulin P, Chiron D, Gavard J & Bidere N (2019) Pannexin-1 limits the production of proinflammatory cytokines during necroptosis. EMBO Rep 20, e47840, doi: 10.15252/embr.201947840.

20. Qu Y, Misaghi S, Newton K, Gilmour LL, Louie S, Cupp JE, Dubyak GR, Hackos D & Dixit VM (2011) Pannexin-1 Is Required for ATP Release during Apoptosis but Not for Inflammasome Activation. The Journal of Immunology 186, 6553–6561, doi: 10.4049/jimmunol.1100478.

21. Woehrle T, Yip L, Elkhal A, Sumi Y, Chen Y, Yao Y, Insel PA & Junger WG (2010) Pannexin-1 hemichannel-mediated ATP release together with P2X1 and P2X4 receptors regulate T-cell activation at the immune synapse. Blood 116, 3475–3484, doi: 10.1182/blood-2010-04-277707.

22. Woehrle T, Yip L, Manohar M, Sumi Y, Yao Y, Chen Y & Junger WG (2010) Hypertonic stress regulates T cell function via pannexin-1 hemichannels and P2X receptors. J Leukoc Biol 88, 1181–1189, doi: 10.1189/jlb.0410211.

23. Chen W, Zhu S, Wang Y, Li J, Qiang X, Zhao X, Yang H, D’Angelo J, Becker L, Wang P, Tracey KJ & Wang H (2019) Enhanced Macrophage Pannexin 1 Expression and Hemichannel Activation Exacerbates Lethal Experimental Sepsis. Sci Rep 9, 160, doi: 10.1038/s41598-018-37232-z.

24. Velasquez S, Malik S, Lutz SE, Scemes E & Eugenin EA (2016) Pannexin1 Channels Are Required for Chemokine-Mediated Migration of CD4+ T Lymphocytes: Role in Inflammation and Experimental Autoimmune Encephalomyelitis. J Immunol 196, 4338–4347, doi: 10.4049/jimmunol.1502440.

25. Penuela S, Kelly JJ, Churko JM, Barr KJ, Berger AC & Laird DW (2014) Panx1 regulates cellular properties of keratinocytes and dermal fibroblasts in skin development and wound healing. J Invest Dermatol 134, 2026–2035, doi: 10.1038/jid.2014.86.

26. Damsky WE, Curley DP, Santhanakrishnan M, Rosenbaum LE, Platt JT, Gould Rothberg BE, Taketo MM, Dankort D, Rimm DL, McMahon M & Bosenberg M (2011) beta-catenin signaling controls metastasis in Braf-activated Pten-deficient melanomas. Cancer Cell 20, 741–754, doi: 10.1016/j.ccr.2011.10.030.

27. Dankort D, Curley DP, Cartlidge RA, Nelson B, Karnezis AN, Damsky WE, Jr., You MJ, DePinho RA, McMahon M & Bosenberg M (2009) Braf(V600E) cooperates with Pten loss to induce metastatic melanoma. Nat Genet 41, 544–552, doi: 10.1038/ng.356.

28. Hooijkaas AI, Gadiot J, van der Valk M, Mooi WJ & Blank CU (2012) Targeting BRAFV600E in an inducible murine model of melanoma. Am J Pathol 181, 785–794, doi: 10.1016/j.ajpath.2012.06.002.

29. Lamprecht MR, Sabatini DM & Carpenter AE (2007) CellProfiler: free, versatile software for automated biological image analysis. Biotechniques 42, 71–75, doi: 10.2144/000112257.

30. Faul F, Erdfelder E, Lang AG & Buchner A (2007) G*Power 3: a flexible statistical power analysis program for the social, behavioral, and biomedical sciences. Behav Res Methods 39, 175–191, doi: 10.3758/bf03193146.

31. Box GEP & Cox DR (1964) An Analysis of Transformations. Journal of the Royal Statistical Society: Series B (Methodological) 26, 211–243, doi: 10.1111/j.2517-6161.1964.tb00553.x.

32. Sviatoha V, Tani E, Kleina R, Sperga M & Skoog L (2010) Immunohistochemical analysis of the S100A1, S100B, CD44 and Bcl-2 antigens and the rate of cell proliferation assessed by Ki-67 antibody in benign and malignant melanocytic tumours. Melanoma Res 20, 118–125, doi: 10.1097/CMR.0b013e3283350554.

33. Xiong TF, Pan FQ & Li D (2019) Expression and clinical significance of S100 family genes in patients with melanoma. Melanoma Res 29, 23–29, doi: 10.1097/CMR.0000000000000512.

34. Ubellacker JM, Tasdogan A, Ramesh V, Shen B, Mitchell EC, Martin-Sandoval MS, Gu Z, McCormick ML, Durham AB, Spitz DR, Zhao Z, Mathews TP & Morrison SJ (2020) Lymph protects metastasizing melanoma cells from ferroptosis. Nature 585, 113–118, doi: 10.1038/s41586-020-2623-z.

35. Nguyen AV & Soulika AM (2019) The Dynamics of the Skin’s Immune System. Int J Mol Sci 20, doi: 10.3390/ijms20081811.

36. Richmond JM & Harris JE (2014) Immunology and skin in health and disease. Cold Spring Harb Perspect Med 4, a015339, doi: 10.1101/cshperspect.a015339.

37. Park SL, Gebhardt T & Mackay LK (2019) Tissue-Resident Memory T Cells in Cancer Immunosurveillance. Trends Immunol 40, 735–747, doi: 10.1016/j.it.2019.06.002.

38. Zhang Z, Liu S, Zhang B, Qiao L, Zhang Y & Zhang Y (2020) T Cell Dysfunction and Exhaustion in Cancer. Front Cell Dev Biol 8, 17, doi: 10.3389/fcell.2020.00017.

39. Freeman GJ, Long AJ, Iwai Y, Bourque K, Chernova T, Nishimura H, Fitz LJ, Malenkovich N, Okazaki T, Byrne MC, Horton HF, Fouser L, Carter L, Ling V, Bowman MR, Carreno BM, Collins M, Wood CR & Honjo T (2000) Engagement of the PD-1 immunoinhibitory receptor by a novel B7 family member leads to negative regulation of lymphocyte activation. J Exp Med 192, 1027–1034, doi: 10.1084/jem.192.7.1027.

40. Jiang JX & Penuela S (2016) Connexin and pannexin channels in cancer. BMC Cell Biol 17 Suppl 1, 12, doi: 10.1186/s12860-016-0094-8.

41. Melnikova VO, Bolshakov SV, Walker C & Ananthaswamy HN (2004) Genomic alterations in spontaneous and carcinogen-induced murine melanoma cell lines. Oncogene 23, 2347–2356, doi: 10.1038/sj.onc.1207405.

42. Karreth FA, Tay Y, Perna D, Ala U, Tan SM, Rust AG, DeNicola G, Webster KA, Weiss D, Perez-Mancera PA, Krauthammer M, Halaban R, Provero P, Adams DJ, Tuveson DA & Pandolfi PP (2011) In vivo identification of tumor-suppressive PTEN ceRNAs in an oncogenic BRAF-induced mouse model of melanoma. Cell 147, 382–395, doi: 10.1016/j.cell.2011.09.032.

43. Meeth K, Wang JX, Micevic G, Damsky W & Bosenberg MW (2016) The YUMM lines: a series of congenic mouse melanoma cell lines with defined genetic alterations. Pigment Cell Melanoma Res 29, 590–597, doi: 10.1111/pcmr.12498.

44. Hooijkaas A, Gadiot J, Morrow M, Stewart R, Schumacher T & Blank CU (2012) Selective BRAF inhibition decreases tumor-resident lymphocyte frequencies in a mouse model of human melanoma. Oncoimmunology 1, 609–617, doi: 10.4161/onci.20226.

45. Zhai Y, Haresi AJ, Huang L & Lang D (2020) Differences in tumor initiation and progression of melanoma in the Braf(CA) ;Tyr-CreERT2;Pten(f/f) model between male and female mice. Pigment Cell Melanoma Res 33, 119–121, doi: 10.1111/pcmr.12821.

46. Freitas-Andrade M, Bechberger JF, MacVicar BA, Viau V & Naus CC (2017) Pannexin1 knockout and blockade reduces ischemic stroke injury in female, but not in male mice. Oncotarget 8, 36973–36983, doi: 10.18632/oncotarget.16937.

47. Aquilino MS, Whyte-Fagundes P, Lukewich MK, Zhang L, Bardakjian BL, Zoidl GR & Carlen PL (2020) Pannexin-1 Deficiency Decreases Epileptic Activity in Mice. Int J Mol Sci 21, doi: 10.3390/ijms21207510.

48. Dakup PP, Porter KI, Little AA, Zhang H & Gaddameedhi S (2020) Sex differences in the association between tumor growth and T cell response in a melanoma mouse model. Cancer Immunol Immunother 69, 2157–2162, doi: 10.1007/s00262-020-02643-3.

49. Ziani L, Safta-Saadoun TB, Gourbeix J, Cavalcanti A, Robert C, Favre G, Chouaib S & Thiery J (2017) Melanoma-associated fibroblasts decrease tumor cell susceptibility to NK cell-mediated killing through matrix-metalloproteinases secretion. Oncotarget 8, 19780–19794, doi: 10.18632/oncotarget.15540.

50. Das A, Ghose A, Naicker K, Sanchez E, Chargari C, Rassy E & Boussios S (2023) Advances in adoptive T-cell therapy for metastatic melanoma. Curr Res Transl Med 71, 103404, doi: 10.1016/j.retram.2023.103404.

51. Hosseini R, Sarvnaz H, Arabpour M, Ramshe SM, Asef-Kabiri L, Yousefi H, Akbari ME & Eskandari N (2022) Cancer exosomes and natural killer cells dysfunction: biological roles, clinical significance and implications for immunotherapy. Mol Cancer 21, 15, doi: 10.1186/s12943-021-01492-7.

52. Cibrian D & Sanchez-Madrid F (2017) CD69: from activation marker to metabolic gatekeeper. Eur J Immunol 47, 946–953, doi: 10.1002/eji.201646837.

53. Edwards J, Wilmott JS, Madore J, Gide TN, Quek C, Tasker A, Ferguson A, Chen J, Hewavisenti R, Hersey P, Gebhardt T, Weninger W, Britton WJ, Saw RPM, Thompson JF, Menzies AM, Long GV, Scolyer RA & Palendira U (2018) CD103(+) Tumor-Resident CD8(+) T Cells Are Associated with Improved Survival in Immunotherapy-Naive Melanoma Patients and Expand Significantly During Anti-PD-1 Treatment. Clin Cancer Res 24, 3036–3045, doi: 10.1158/1078-0432.CCR-17-2257.

54. Varela-Vazquez A, Guitian-Caamano A, Carpintero-Fernandez P, Fonseca E, Sayedyahossein S, Aasen T, Penuela S & Mayan MD (2020) Emerging functions and clinical prospects of connexins and pannexins in melanoma. Biochim Biophys Acta Rev Cancer 1874, 188380, doi: 10.1016/j.bbcan.2020.188380.

55. Tengesdal IW, Dinarello A, Powers NE, Burchill MA, Joosten LAB, Marchetti C & Dinarello CA (2021) Tumor NLRP3-Derived IL-1β Drives the IL-6/STAT3 Axis Resulting in Sustained MDSC-Mediated Immunosuppression. Frontiers in Immunology 12, Original Research, doi: 10.3389/fimmu.2021.661323.

56. Lelliott EJ, Cullinane C, Martin CA, Walker R, Ramsbottom KM, Souza-Fonseca-Guimaraes F, Abuhammad S, Michie J, Kirby L, Young RJ, Slater A, Lau P, Meeth K, Oliaro J, Haynes N, McArthur GA & Sheppard KE (2019) A novel immunogenic mouse model of melanoma for the preclinical assessment of combination targeted and immune-based therapy. Sci Rep 9, 1225, doi: 10.1038/s41598-018-37883-y.

57. Peng W, Chen JQ, Liu C, Malu S, Creasy C, Tetzlaff MT, Xu C, McKenzie JA, Zhang C, Liang X, Williams LJ, Deng W, Chen G, Mbofung R, Lazar AJ, Torres-Cabala CA, Cooper ZA, Chen PL, Tieu TN, Spranger S, Yu X, Bernatchez C, Forget MA, Haymaker C, Amaria R, McQuade JL, Glitza IC, Cascone T, Li HS, Kwong LN, Heffernan TP, Hu J, Bassett RL, Jr., Bosenberg MW, Woodman SE, Overwijk WW, Lizee G, Roszik J, Gajewski TF, Wargo JA, Gershenwald JE, Radvanyi L, Davies MA & Hwu P (2016) Loss of PTEN Promotes Resistance to T Cell-Mediated Immunotherapy. Cancer Discov 6, 202–216, doi: 10.1158/2159-8290.CD-15-0283.

58. Lin Z, Huang L, Li SL, Gu J, Cui X & Zhou Y (2021) PTEN loss correlates with T cell exclusion across human cancers. BMC Cancer 21, 429, doi: 10.1186/s12885-021-08114-x.

59. Massi D, Rulli E, Cossa M, Valeri B, Rodolfo M, Merelli B, De Logu F, Nassini R, Del Vecchio M, Di Guardo L, De Penni R, Guida M, Sileni VC, Di Giacomo AM, Tucci M, Occelli M, Portelli F, Vallacchi V, Consoli F, Quaglino P, Queirolo P, Baroni G, Carnevale-Schianca F, Cattaneo L, Minisini A, Palmieri G, Rivoltini L, Mandala M & Italian Melanoma I (2019) The density and spatial tissue distribution of CD8(+) and CD163(+) immune cells predict response and outcome in melanoma patients receiving MAPK inhibitors. J Immunother Cancer 7, 308, doi: 10.1186/s40425-019-0797-4.

