## Supplementary figures and images for "Global Pannexin 1-deletion increases tumor-infiltrating lymphocytes in the BRAF/Pten mouse melanoma model"

### Supplemental Figure 1

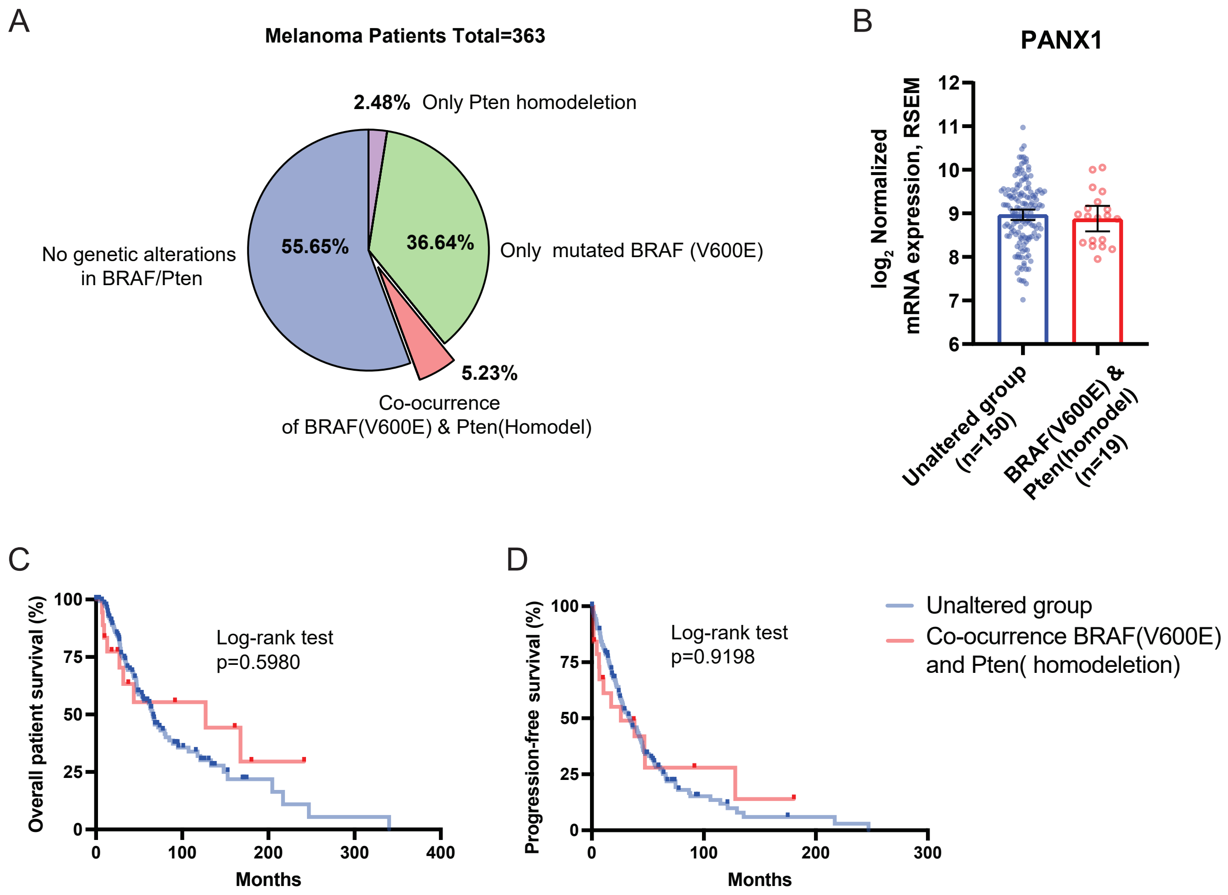
